# Money or smiles: Independent ERP effects of associated monetary reward and happy faces

**DOI:** 10.1101/325829

**Authors:** Wiebke Hammerschmidt, Louisa Kulke, Christina BrÖring, Annekathrin Schacht

## Abstract

In comparison to neutral faces, facial expressions of emotion are known to gain attentional prioritization, mainly demonstrated by means of event-related potentials (ERPs). Recent evidence indicated that such a preferential processing can also be elicited by neutral faces when associated with increased motivational salience via reward. It remains, however, an open question whether impacts of inherent emotional salience and associated motivational salience might be integrated. To this aim, expressions and outcomes were orthogonally combined. Participants (*N*=42) learned to explicitly categorize happy and neutral faces as either reward- or zero-outcome-related via an associative learning paradigm. ERP components (P1, N170, EPN, and LPC) were measured throughout the experiment, and separately analyzed before (learning phase) and after (consolidation phase) reaching a pre-defined learning criterion. Happy facial expressions boosted early processing stages, as reflected in enhanced amplitudes of the N170 and EPN, both during learning and consolidation. In contrast, effects of monetary reward became evident only after successful learning and in form of enlarged amplitudes of the LPC, a component linked to higher-order evaluations. Interactions between expressions and associated outcome were absent in all ERP components of interest. The present study provides novel evidence *that* acquired salience impacts stimulus processing but independent of the effects driven by happy facial expressions.

## Introduction

Because of limited sensory and cognitive resources, the human brain has evolved efficient selection mechanisms that bias perception in favour of salient, i.e. behaviourally relevant or physically distinct, information. Stimuli of increased salience have been demonstrated to directly capture attention and impact visual processing capacities [e.g., 1,2], resulting in facilitated sensory encoding even at initial processing stages [e.g., 3]. Faces, and in particular facial expressions of emotion, were demonstrated to convey particularly high salience, as they not only provide important information about others in social interactions, but also have an intrinsic relevance to assure survival and well-being. Therefore, it has been assumed that humans have evolved a biological preparedness to rapidly detect emotional expressions [e.g., 4]. For facial expressions of emotion, preferential processing has been unveiled both at the behavioural and neural level, and in particular for angry facial expressions [e.g., 5–9]. However, for happy facial expressions faster and more accurate recognition compared to other expressions has been demonstrated [10,11], that is potentially based one the exclusive role of happiness as a positive expression [12,13]. In addition, as humans are highly social beings [e.g., 14], facial expressions of emotion are not only emotionally but also motivationally relevant, as, for instance, a happy face might carry a rewarding value similar to other reinforcers [e.g., 15,16]. Traditional theories of attention focused on bottom-up (i.e., stimulus-driven) and top-down (i.e., goal-directed) attention mechanisms [e.g., 17] to explain how relevant stimuli are preferentially processed. However, such accounts have recently been challenged by studies demonstrating a preferential processing of previously reward-associated stimuli, which occurs even when the stimuli themselves do not carry increased salience, when they are task-irrelevant, or when the reward is suspended over time [18]. Hence, Anderson proposed a general value-driven attention mechanism to explain the attentional prioritization of not only stimuli of inherent emotional salience but also of stimuli that acquired their salience through learning processes. Supporting evidence for this assumption comes from findings indicating overlapping neural activity in the ventromedial prefrontal cortex (vmPFC) and the ventral striatum, elicited by both emotional facial expressions and monetary reward [19]. Furthermore, motivational relevance has been widely equated with emotional stimulus valence or seen as a precursor of emotional significance in some scientific approaches [e.g., 20,21].

An excellent tool to gain insights into the neuro-cognitive mechanisms underlying the prioritized processing of emotional stimuli are event-related brain potentials (ERPs) since they allow dissociating between different processing stages. In the domain of facial expressions of emotion, a large number of studies revealed rather robust modulations of dissociable ERP components over time: The Early Posterior Negativity (EPN), a typical emotion-related ERP component reflecting an enhanced sensory encoding of stimuli carrying inherent salience, occurs around 150-200 ms after face stimulus onsets [6,8,9] and has been demonstrated to be elicited by emotional expressions, including happiness/joy [22–25]. The EPN to emotional stimuli is typically followed by the Late Positivity Complex (LPC), developing around 300 ms and lasting for several hundred milliseconds [e.g., 7]. The LPC – linked to higher-order processes of stimu-lus evaluation and likewise termed as Late Positive Potential [e.g., 9] – has been shown to be modulated by angry expressions, presumably due to their particular evolutionary relevance [9]. However, also happy expressions amplified the LPC component [6,8,24]. Moreover, the P1 component, peaking around 100 ms at occipital electrodes, presumably reflects rapid activation of the extrastriate visual cortex [26] and is mainly impacted by negative expressions [e.g., 8,25,27]. The N170 component, typically following the P1 in face processing, is an occipito-temporal negativity linked to holistic face perception [28]. Several studies reported N170 modulations by emotional expressions, including happy expressions [e.g., 24,29], but also negative facial expressions elicited enhanced N170 modulations [for reviews, see 30,31].

In addition to these robust ERP effects elicited by facial expressions of emotions, also neutral faces associated with motivational salience were reported to impact dissociable ERP components over time. A recent study by Hammerschmidt and coworkers [25] directly compared the neural correlates of processing facial expressions of emotion and neutral faces associated with motivational salience by means of ERPs. Interestingly, reward-associated neutral faces elicited enhanced amplitudes of the P1 component, similar to P1 amplification by angry facial expressions. Whereas the prioritization of associated motivational salience was restricted to initial processing stages (P1), beneficial processing of facial expressions of emotions spread over to subsequent stages of more elaborative stimulus processing (EPN, LPC). In other studies, however, neutral faces implicitly associated with monetary reward have been shown to elicit enhanced LPC amplitudes [32], replicating previous findings that the LPC component seems to be sensitive to reward associations [33,34]. According to the value-driven attention mechanism [18], the processing of inherent emotional and associated motivational salience should share certain similarities. However, evidence is inconclusive with regard to this assumption, as the majority of previous studies has investigated these types of salience separate from each other. Interactions of associated reward and emotional expression were reported on reaction times [35], but only when the facial expression was task-relevant. Further, Yao and colleagues [36] demonstrated that the preferential processing of angry expressions can be extenuated through reward associations. However, the authors only investigated effects on the N2pc component, linked to spatial attention [37], and disregarded the investigation of emotion-related ERP components.

The present study aimed at clarifying whether the preferential processing of inherently happy facial expressions might be further boosted by acquired motivational salience (reward). To this aim, we used an explicit associative learning paradigm similar to the study by Hammerschmidt and co-workers [25]. Considering that in this previous study effects of associated salience were restricted to the reward condition, here only happy and neutral faces were orthogonally associated with monetary gain or zero outcome, respectively. Directly after participants reached a predefined learning criterion, a consolidation phase was added to strengthen the learned associations. ERPs were recorded to compare the effects of the factors expressions, outcome and their potential interaction over different stages of face processing.

Replicating previous findings, we hypothesized reward associations to be learned faster than zero-outcome-associations [25,34,38], as well as faster reaction times for reward compared to zero-outcome-associations [25]. In line with the literature, happy faces were expected to trigger the typical emotion-related EPN component [e.g., 8,22,25,32,40] both in the learning and consolidation phase. A reward-modulation on the P1 component was expected for neutral faces associated with monetary gain [25]. In addition, if associated motivational and inherent emotional salience would be integrated at sensory processing stages, happy faces associated with reward were expected to elicit even stronger P1 modulations than neutral faces associated with monetary gain. The potential interaction of emotional expression and associated outcome was investigated on all measurements, behavioral data and ERPs.

Finally, one might expect an improvement of participants’ mood from the beginning to the end of the experimental session for several reasons, that is, the absence of negative stimuli throughout the experiment, the increase of monetary gain during successful learning, and the general release of having an experiment completed. We therefore quantified pre- and post-experimental mood state of the participants on multiple dimensions (mood, calmness, alertness), in order to detect and exclude possible outliers from the analyses.

## Materials and Methods

### Participants

Data was collected from 48 participants. Four participants were excluded, as they did not reach the required learning criterion within 10 to 30 blocks, and two participants due to excessive artifacts during EEG recordings. The remaining forty-two participants (20 female) had an age range between 19 and 30 years (mean age = 23.9 years, *SD* = 2.7), normal or corrected-to-normal vision and no neurological or psychiatric disorders according to self-report. All participants were right-handed [according to 41] and were reimbursed with their individual bonus, ranging between 38.10 and 59.70 euro (*M* = 49.82 euro, *SD* = 5.47 euro).

### Stimuli

Sixteen colored facial stimuli (8 female, 8 male) were selected from the Karolinska Directed Emotional Faces (KDEF) database [41], showing happy and neutral expressions respectively. A grey ellipsoid mask, ensuring a uniform figure/ground contrast, surrounded the stimuli within an area of 130 x 200 pixels (4.59 x 7.06 cm) and let only the internal face area visible. Facial stimuli were matched for luminance across conditions (according to Adobe Photoshop CS6^TM^), *F*(1,30) = 2.907, *p* =0.099, and were presented at a central position on the screen on a light gray background, corresponding to a visual angle of 4.6° x 7.1°. Feedback symbols were presented in grey circles in the center of the screen (248 x 248 pixels, 5 x 5 cm) and were matched in contrast, luminance, and visual complexity (800 pixels respectively): a green plus (correct reward condition), a dark grey equality sign (correct zero-outcome condition) or a red cross (error) (please see Fig 1).

**Figure 1.**
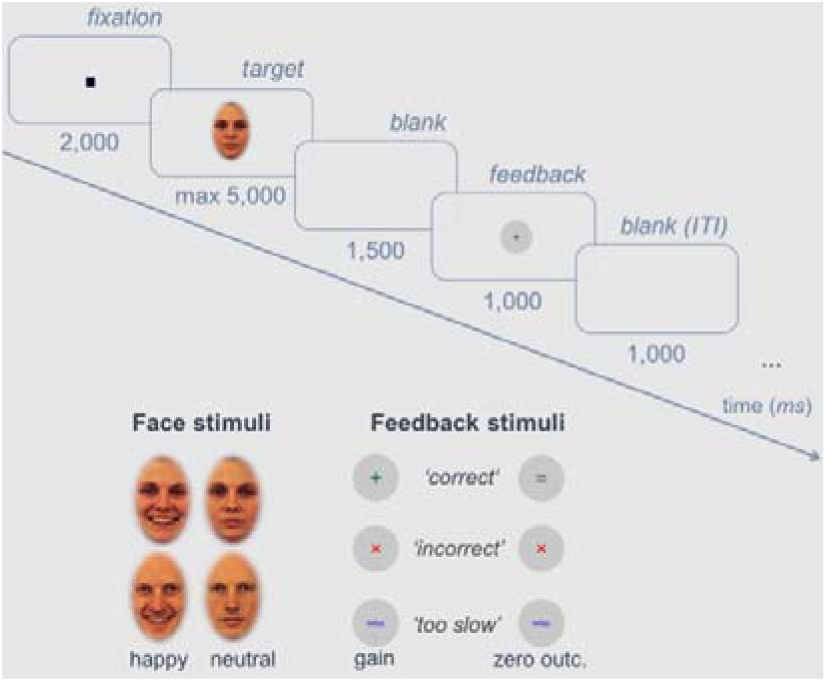
Experimental Procedure. Trial scheme with detailed time sequence of the associative learning task and exemplary face and feedback stimuli.

### Procedure

The study was conducted in accordance with the Declaration of Helsinki and approved by the local Ethics committee of the Institute of Psychology at the University of Goettingen. Participants were informed about the procedure and gave written informed consent prior to the experiment. Participants were seated in a dimly lit, sound-attenuated room at a viewing distance of 57 cm to the computer screen. A chin rest was used.

During the experiment, eight inherently happy and eight neutral faces were associated with monetary gain or zero-outcome via an associative learning paradigm [similar to 25]. The participants’ task was to learn the correct outcome-expression assignment for each of the faces presented. As no test trials were provided, the first block had to be answered by chance. The feedback scheme was explained prior to the experiment: In case of correct classifications, faces that had to be categorized as reward-related were associated with +20 cents, and faces that had to be categorized as zero-outcome-related were associated with 0 cents. For incorrect classifications, participants lost −10 cents, independent of the outcome category. If the participants missed to answer a trial within 5000 ms, 50 cents were removed from their bonus. Responses were given by button press with two fingers of the dominant hand; response-to-button assignment was balanced across participants, as well as face-to-expression/outcome assignment, but re-mained the same for each participant. Stimuli were presented in 40 blocks; each block consisted of all sixteen facial stimuli, presented in fully randomized order within a given block. The experiment thus consisted of 640 trials in total. Blocks were separated by a self-determined break where information about the current amount of the individual bonus was provided. A learning criterion was defined (48 of the last 50 trials correct) to assure successful learning. If the learning criterion was not reached within 10 to 30 blocks, data was excluded from analysis (*N* = 4). After meeting the learning criterion, the remaining trials, up to 40 blocks in total, were presented subsequently (consolidation phase). During both learning and consolidation phase, the trial scheme was as follows (see Fig 1): A black fixation point (5 x 5 pixels) was presented for 2000 ms in each trial, followed by the face for a maximum of 5000 ms, disappearing with button press. Afterwards, a blank screen for 1500 ms and the feedback symbol for 1000 ms were presented; the inter-trial interval (blank) was 1000 ms. In order to obtain potential changes in the mood of participants, the German Multidimensional Mood Questionnaire [MDBF; 43] was completed before and after the experimental task.

### EEG recording, pre-processing and analyses

The electroencephalogram (EEG) was recorded from 64 electrodes, placed in an electrode cap (Easy-Cap, Biosemi, Amsterdam, The Netherlands) according to the extended 10-20 system [43]. The common mode sense (CMS) electrode and the driven right leg (DRL) passive electrode were used as reference and ground electrodes (cf., http://www.biosemi.com/faq/cms&drl.htm)

Six external electrodes were used, inferior and laterally to the eyes to record blinks, and on the left and right mastoids. Signals were recorded at a sampling rate of 2048 Hz (downsampled to 500 Hz for ERP analysis) and a bandwidth of 104 Hz (http://www.biosemi.com/faq/adjust_filter.htm), offline filtered with a Low Cutoff (0.03183099 Hz, Time constant 5 s, 12 dB/oct), a High Cutoff (40 Hz, 48 dB/oct), and a Notch Filter (50 Hz). Data was processed with BrainVision Analyzer (Brain Products GmbH, Munich, Germany), average-referenced and corrected for blinks using Surrogate Multiple Source Eye Correction with default parameters [MSEC; 45, application is detailed in 46] as implemented in BESA (Brain Electric Source Analysis, MEGIS Software GmbH, Gräfelfing, Germany). The continuous EEG signal was segmented into epochs of 1200 ms, starting 200 ms before face stimulus onset and referring to a 200 ms prestimulus baseline. Electrodes with noisy or no signal were interpolated by spherical splines in BrainVision Analyzer (Order of splines: 4; maximal degree of Legendre Polynomials: 10; Lambda: 1E-05). Epochs containing artifacts (criteria: voltage steps > 50 μV, 200μV/200 ms intervals difference of values, amplitudes exceeding - 150μV/150 μV, activity < 0.5 μV) were eliminated. Segments were averaged per Subject, Phase (2 – learning, consolidation), Expression (2 – happy, neutral) and Outcome (2 – reward, zero outcome). Based on a previous study [25], time windows and regions of interest (ROIs) electrodes for target face-related ERP components were chosen as follows: i) P1: 75-125 ms, O1 and O2; ii) N170: 130-180 ms, P9 and P10; iii) EPN: 200-350 ms, P9, P10, Iz, Oz, O1, O2, PO7 and PO8; iv) LPC: 350-700 ms, Pz, P1, P2, CPz and POz. The P1 component was quantified as the most positive peak (with O2 as reference electrode), the N170 component as the most negative peak (with P10 as reference electrode), and EPN and LPC were quantified as mean amplitudes across electrodes. For statistical analysis, repeated-measures (rm)ANOVAs were computed, including the factors Expression (2 – happy, neutral) and Outcome (2 – reward, zero outcome), separate for the learning and consolidation phase.

### Analyses of behavioral data

To investigate potential differences in learning curves between conditions, posterior distributions for the probability (the coefficient p of a Bernoulli distribution) to attribute the outcome category correctly were modeled. The number of trials until the learning criterion was met differed between participants. To account for these differences in trial numbers, the proportion of time (until the learning criterion was met) was considered (see Fig 2). Significant differences between the learning curves were defined based on a criterion of non-overlapping 99 % simultaneous credible bands [for more details, see 25].

**Figure 2.**
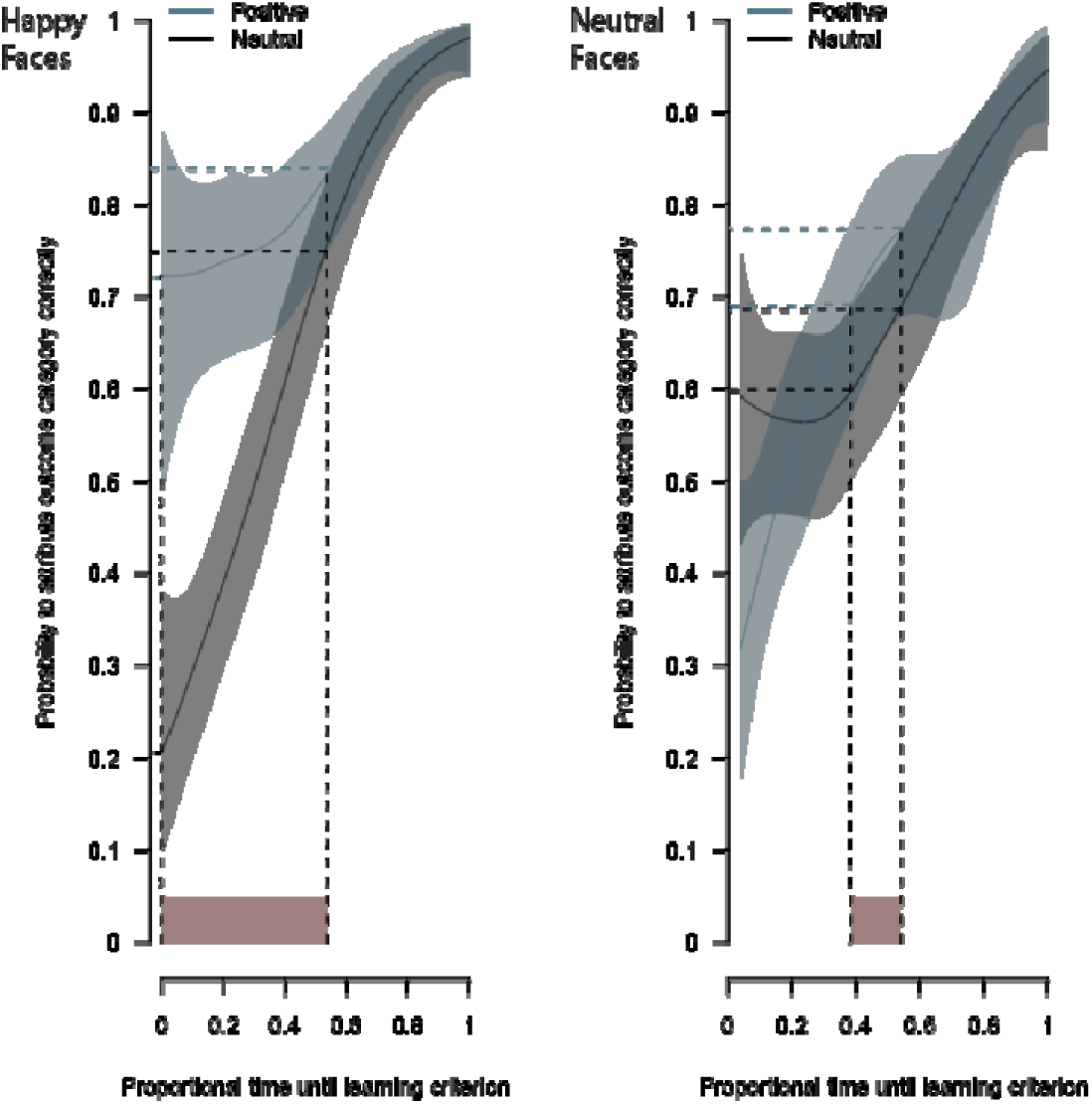
Learning curves. Posteriori mean probabilities to attribute the outcome category correctly during the learning phase (illustrated by horizontal dashed lines) at the lower and upper bounds of the time intervals until the learning criterion was met (illustrated by red areas)..

For accuracy data in the consolidation phase and reaction times in both phases, repeated-measures (rm)ANOVAs were computed, including the factors Expression (2 – happy, neutral) and Outcome (2 – reward, zero outcome). Outliers were identified as reaction times (RTs) below 200 ms or exceeding +2*SDs* from the mean per condition and were excluded from behavioral data analysis.

## Results

### Learning Phase

#### Behavioral Data

Participants (*N* = 42) learned the outcome associations adequately within 10 to 26 blocks (*M* = 17.2 blocks, *SD* = 4.8 blocks). In the happy face condition, reward associations were learned faster, differing from zero-outcome associations from the beginning until 53.9% of the learning criterion was met. At this time, participants were correct 74.9% of the time for zero-outcome associations and 83.9% for reward associations. In the neutral face condition, reward associations were learned faster, differing from zero-outcome associations from 38.5% until 54.3% of the learning criterion was met. Note, however, that in the very beginning, there was an advantage for zero-outcome associations compared to reward associations until 13.3% of the learning criterion was met (see Fig 2).

Reaction times (RTs) were shorter for happy compared to neutral expressions, *F*(1,41) = 7.393, *p* = 0.010, 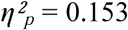, as well as for reward-compared to zero-outcome associated faces, *F*(1,41) = 10.301, *p* = 0.003, 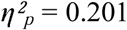. An interaction between Expression and Outcome was absent. Mean RTs per experimental condition are summarized in Table 1.

**Table 1.** Mean reaction times in ms and accuracy in % for learning and consolidation phase (SEMs in parentheses), contrasted for all factor levels of Expression and Outcome.

|  | Emotion | Outcome | RTs | Accuracy* |
| --- | --- | --- | --- | --- |
| Learning Phase | Happy | Reward | 1398 (44) | - |
|  |  | Zero Outcome | 1477 (40) | - |
|  | Neutral | Reward | 1480 (41) | - |
|  |  | Zero Outcome | 1530 (42) | - |
| Consolidation Phase | Happy | Reward | 942 (27) | 99.3 (0.2) |
|  |  | Zero Outcome | 1006 (29) | 99.1 (0.2) |
|  | Neutral | Reward | 986 (25) | 98.8 (0.3) |
|  |  | Zero Outcome | 1016 (28) | 99.1 (0.3) |

\*Adequate accuracy of each participant during the learning phase was assured by a required learning criterion (48 of the last 50 trials correct).

Reaction times (RTs) revealed a main effect of the factor Emotion, *F*(1,41) = 5.647, *p* = 0.022, 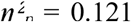, with faster reaction times for happy compared to neutral facial expressions, and the factor Outcome, *F*(1,41) = 11.347, *p* = 0.002, 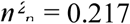, with faster reaction times for reward-in comparison to zero-outcome-associated faces; an interaction effect was absent. Mean reaction times per experimental condition are summarized in Table 1.

#### ERPs to Target Faces

Significant main effects of the factor Expression were revealed on the N170 component, *F*(1,41) = 14.855, *p* < 0.001, 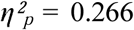, and on the EPN component, *F*(1,41) = 42.405, *p* < 0.001, 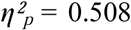, both reflecting larger negativities for happy in comparison to neutral expressions (see Fig 3). No effect of the factor Outcome occurred. P1 and LPC were not modulated by the factors Expression or Outcome. There was further no evidence for interactions between the factors on all ERP components of interest.

**Figure 3.**
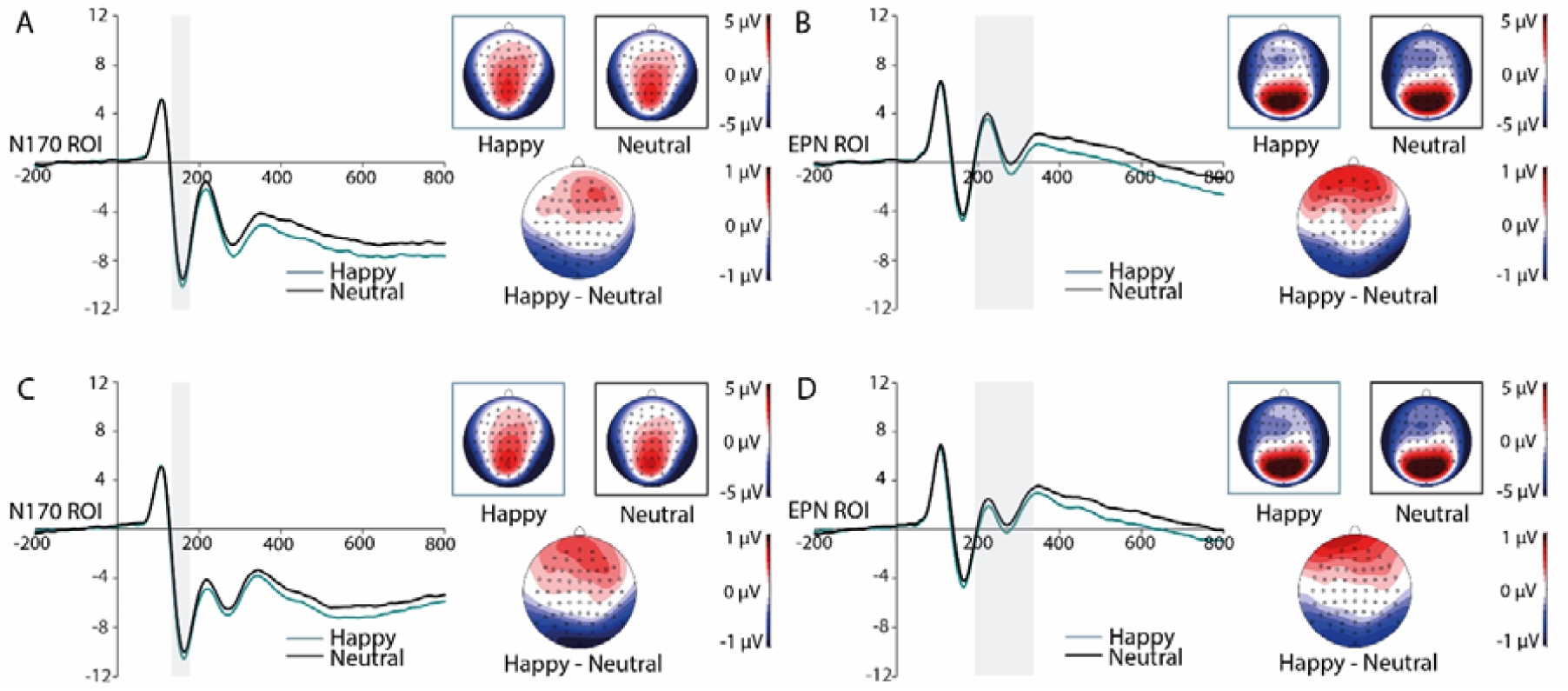
Grand-averaged ERPs at N170-ROI electrodes for happy and neutral faces during the learning (A) and consolidation phase (C) with corresponding scalp distributions and to-pographies of ERP differences between indicated emotion categories. Grand-averaged ERPs at EPN-ROI electrodes for happy and neutral faces during the learning (B) and consolidation phase (D) with corresponding scalp distributions and topographies of ERP differences between indicated emotion categories. Highlighted areas display the time windows of ERP analyses.

### Consolidation Phase

#### Behavioral Data

Accuracy was at ceiling (> 99%) for all conditions (see Table 1) during the consolidation phase and did not differ in terms of the factors Expression and Outcome, *F*(1,41) < 1. Analyses on RTs revealed a main effect of the Factor Outcome, *F*(1,41)= 17.235, *p* < 0.001, 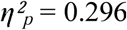, with faster RTs for reward-compared to zero-outcome associated faces. Neither the main effect of the factor Expression nor the interaction between both factors reached significance. Mean RTs and accuracy values per experimental condition are summarized in Table 1.

#### ERPs to Target Faces

Significant main effects of the factor Expression were obtained on the N170, *F*(1,41) = 21.015, *p* < 0.001, 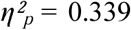, and on the EPN component, *F*(1,41) = 15.923, *p* < 0.001, 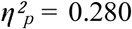, both with enhanced negativities for happy compared to neutral facial expressions (see Fig 3). ERP modulations by the factor Outcome were restricted to the LPC component, *F*(1,41) = 5.260, *p* = 0.027,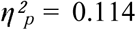, with increased amplitudes for reward-compared to zero-outcome-associated faces, (see Fig 4). The P1 component was not modulated by factors Expression and Outcome. Interactions between the factors Expression and Outcome were absent on all components of interest.

**Figure 4.**
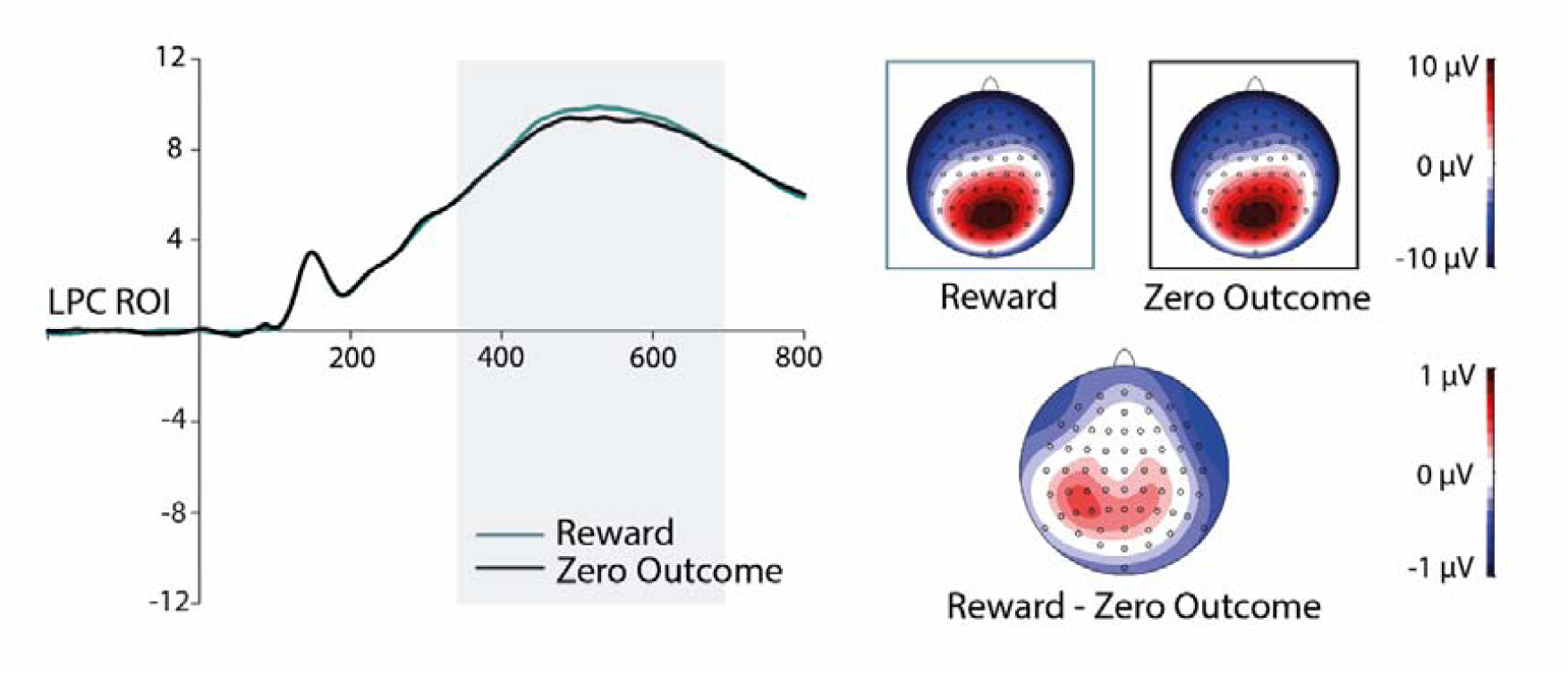
Grand-averaged ERPs at LPC-ROI electrodes in response to reward- and neutral-associated faces during the consolation phase with corresponding scalp distributions (left panel) and topographies of ERP differences (right panel) between indicated motivation categories. Highlighted area displays the time windows of ERP analysis.

As a potential interaction of Expression and Outcome on LPC amplitudes in the consolidation phase was of particular interest, it was followed up with a Bayesian analysis using the “BayesFactor” Package [46] in R [47] with Cauchy priors based on [48]. The “anovaBF” function was used to predict the LPC amplitude averaged across the electrode cluster from Outcome (reward, zero outcome), Expression (happy, neutral) and the interaction term. A comparison of the Bayes factor for the model including the interaction and main effects and the model including only the main effects showed that the interaction of Outcome and Expression is 1.85 times more likely to have no effect on LPC amplitude than to have an effect, *BF*_10_ = 0.54.

#### Mood

Participants’ mood (according to MDBF) was significantly better after the associative learning task, *F*(1,41) = 9.718, *p* = 0.003, 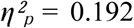, whereas alertness was reduced compared to the beginning of the experiment, *F*(1,41) = 16.034, *p* < 0.001, 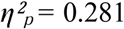. The task did not impact participants’ calmness.

## Discussion

The aim of the present study was the investigation of a potential integration of inherent emotional and associated motivational salience. Such conjunction into an integrated salience appears reasonable, considering the manifold similarities in their processing characteristics [18]. In order to test this assumption, happy and neutral faces were associated with monetary gain or zero outcome via explicit associative learning, successfully implemented in previous research [e.g., 25,32,34]. The experimental data was divided into a learning and consolidation phase, defined by the individual performance of the participants, allowing to investigate ERPs during and after successful learning. On average, outcome associations had no impact on visual processing during the learning phase, whereas in the consolidation phase, LPC amplitudes, referred to an elaborative processing of relevant stimuli, were enhanced by reward associations. The amplified LPC to reward-associated faces, independent of their expressions, replicates studies that implemented implicit reward learning with neutral faces [32] or explicit associations of reward to (neutral) letters from unfamiliar alphabets [33,34]. Importantly, the learning data corroborate our finding of a reward-driven LPC, irrespective of the facial expression. Associations of happy faces with reward were learned the fastest, potentially due to an advantage of congruency of expression and outcome valence. However, neutral faces associated with zero outcome were learned better in the beginning but were outperformed by reward associations over time. Across both phases, an advantageous effect of reward became also evident in reaction times, as responses to reward-associated faces were faster than those to faces associated with zero outcome. Interestingly, the facilitated processing of happy compared to neutral faces, as reflected in shorter reaction times, was limited to the learning phase but vanished during consolidation. Finally, although the task was demanding, as indicated by a decrease of participants’ alertness, mood increased. Together, these findings highlight the increased behavioral relevance of reward associations.

Early P1 modulations were expected to be elicited by reward associations [25]. For their absence, three explanations might be considered: First, a special feature of the present experiment was the restriction to happy and neutral faces as target stimuli on the one hand and gain and zero-outcomes on the other hand, leading to the complete absence of any aversive stimulus. One might assume that effects of reward are stronger or even limited to conditions when a negative counterpart (e.g. angry face or monetary loss) is present. Second, in contrast to our previous study [25], no delay between learning and subsequent testing was implemented, while also during the consolidation phase feedback about monetary outcome was provided. The consolidation of emotional or rather arousing stimuli has been suggested to require time [49], however, also P3 effects modulated by monetary reward were found without overnight consolidation suggesting that this is not mandatory for the occurrence of reward associations [34]. Third, similar studies [33,38,50] used a different task during delayed testing, while in the present study, the categorization task remained the same and the task relevance of the stimuli did not change throughout the experiment. Therefore, the experimental design, e.g. the task, might play a crucial role in understanding impacts of associated motivational salience and need to be spotlighted in further research.

Happy expressions impacted the face-sensitive N170 and the typical emotion-related EPN component both during learning and consolidation. The N170 reflects the configural encoding of a facial stimulus [28]. There is still an ongoing debate whether this process might be impacted by facial expressions of emotion [for reviews, see 30,31]. However, several studies could demonstrate that the N170 component might be modulated by happy facial expressions [e.g., 24,29]. The finding of an EPN modulation by happy facial expressions is in line with the conventional link of the EPN to an enhanced encoding of relevant sensory information [7,8,39] that occurs independent of context and task demands. Interestingly, as can be seen in Fig 3, the difference distributions of the N170 resembled those of the EPN, indicating a potential overlap of these two ERP components [7,39]. Future research is needed to fully dissociate these two prominent ERP components and their potential modulations by emotional aspects.

Importantly, no interaction of the factors Expression and Outcome was detected, neither on the behavioral level nor in any of the ERP components of interest, indicating that no integration of the two sources of salience takes place. This absence of interaction effects corroborates studies on emotional words [51] and behavioral findings for faces in a study, where the emotional expression was not task-relevant [35]. In addition, a decrease of the preferential processing of angry (but not happy) faces by associations with monetary reward was previously demonstrated in modulations of the N2pc [36], a component linked to spatial attention [37]. This finding might further emphasize the relevance of a negative/aversive counterpart to positive stimuli (e.g., happy faces) presented in associative learning paradigms.

## Conclusion

Together, the results of the present study only partially support the value-driven attention mechanism proposed by Anderson [18]. According to his assumption, the prioritized processing of stimuli carrying associated motivational and inherent emotional salience should be highly similar. Such similarity should not only be observed in (facilitated) behavior but moreover in common functional loci that should be reflected in additive or even interactive effects at dissociable ERP components over time. In the present study, we orthogonally combined both types of salience in an associative learning paradigm and measured ERPs throughout the experiment. Importantly, our main finding indicated strong differences that these two types of salience-specific impacts at different processing stages, reflected by the occurrence of their effects in different ERP components/time windows, diverging brain topographies of these ERP modulations, and finally the absence of interaction effects, under the given experimental conditions.

## Acknowledgments

We thank Maren Mette for her contribution to data collection and Holger Sennhenn-Reulen for providing the code for behavioral data analysis.

